# Adaptive current-flow models of ECT: Explaining individual static impedance, dynamic impedance, and brain current density

**DOI:** 10.1101/2020.11.08.373712

**Authors:** Gozde Unal, Jaiti K. Swami, Carliza Canela, Samantha L. Cohen, Niranjan Khadka, Mohammad Rad, Baron Short, Miklos Argyelan, Harold A. Sackeim, Marom Bikson

## Abstract

**Background:** Improvements in electroconvulsive therapy (ECT) outcomes have followed refinement in device electrical output and electrode montage. The physical properties of the ECT stimulus, together with those of the patient’s head, determine the impedances measured by the device and govern current delivery to the brain and ECT outcomes.

**Objective:** However, the precise relations among physical properties of the stimulus, patient head anatomy, and patient-specific impedance to the passage of current are long-standing questions in ECT research and practice.

**Methods:** We developed anatomical MRI-derived models of transcranial electrical stimulation (tES) that included changes in tissue conductivity due to local electrical current flow. These “adaptive” models simulate ECT both during therapeutic stimulation using high (~1 A) current and when dynamic impedance is measured, as well as prior to stimulation when low (~1 mA) current is used to measure static impedance. We modeled two scalp layers: a superficial scalp layer with adaptive conductivity that increases with electric field up to a subject specific maximum 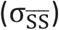, and a deep scalp layer with a subject-specific fixed conductivity (σ_DS_).

**Results:** We demonstrate that variation in these scalp parameters explain clinical data on subject-specific static impedance and dynamic impedance, their imperfect correlation across subjects, their relationships to seizure threshold, and the role of head anatomy. Adaptive tES models demonstrate that current flow changes local tissue conductivity which in turn shapes current delivery to the brain in a manner not accounted for in fixed tissue conductivity models.

**Conclusions:** Our predictions that variation in individual skin properties, rather than other aspects of anatomy, largely govern the relationship between static impedance, dynamic impedance, and current delivery to the brain, are themselves subject to assumptions about tissue properties. Broadly, our novel pipeline for tES models is important in ongoing efforts to optimize devices, personalize interventions, and explain clinical findings.

## Introduction

Ongoing advancements in the efficacy or specificity (reduced adverse effects) of electroconvulsive therapy (ECT) rely on refinement in dose; namely electrode montage and stimulation waveform and intensity [1,2]. Dose optimization has followed heuristic approaches, and controversies remain unreconciled despite decades of research [3,4]. Modern ECT devices deliver current-controlled (800-900 mA) pulses, so that applied voltage is adjusted based on the encountered impedance. Devices report the “dynamic impedance” during the passage of the ECT stimulus. Prior to stimulation, ECT devices report a “static impedance” using low intensity (~1 mA) high-frequency test currents. While static impedance and dynamic impedance has long been recognized as markers of individual difference, their etiology and consequences are undetermined [5] – including how they impact on seizure induction [6–9].

The finding that dynamic impedance is lower than static impedance is consistent with an increase in tissue conductivity in response to high (dynamic impedance) verse low (static impedance) current intensity. But it is unclear which tissue layers are responsible for this impedance difference, how current flow to the brain is then altered, and with what implications to seizure genesis?

While significant covariation between static and dynamic impedance has been reported [10], debate about whether there is a meaningful correlation has spanned decades [5,11,12]. One aim of this study was to explain the imperfect relationship between static and dynamic impedance. We use clinical trial datasets with standardized electrode preparation conditions to reexamine the relations between static and dynamic impedance values under fastidious conditions. In a sample of normal subjects, we systematically manipulated electrode preparation factors (contact area, adherence) to determine the impact of preparation protocol on static impedance across subjects. Novel current-flow models were developed and experientially constrained based on ECT subjects anatomical imaging and impedance data, to systematically explain what factors drive verse limit the correlation between static and dynamic impedance.

For a given ECT electrical dose, the pattern of current delivery to the brain is determined by individual anatomy and electric conductivity of each tissue compartment (e.g. skin, skull) [10,13,14]. In theory, changing tissue conductivity during the passage of the ECT stimulus would influence the distribution of current density in brain. Correlations between dynamic impedance and seizure threshold have been shown [10] but not reliably [8,9]. A further aim of this study, using current-flow models, is to understand how changing tissue conductivity (alongside other anatomical factors) govern brain current flow during ECT and how this is reflected in overall impedance.

We developed an individualized (MRI-derived) finite element method (FEM) to model transcranial electrical stimulation while incorporating “adaptive” changes in tissue conductivity by local current flow. These models explain the source of individual difference in ECT static and dynamic impedance, how they relate, and how they impact seizure threshold. More generally, these state-of-the-art models demonstrate that adaptive change in tissue conductivity shapes current delivery across the head, resulting in different patterns than predicted by fixed-conductivity models.

## Methods

### Electrode preparation on static impedance: controlled experiments in healthy subjects

We conducted a study to evaluate the role of electrode preparation on static impedance using healthy subjects (n=3) within whom six electrode preparation techniques could be compared. All comparisons were made using disposable, adherent pads as electrodes (Thymapads, Somatics, LLC, Venice, FI). The electrode preparation techniques included: 1) Skin cleaning with saline and application of Pretac (Pharmaceutical Innovations, Newark, New Jersey); 2) Skin cleaning with alcohol (99%) and application of Pretac; 3) Skin cleaning with saline and application of Pretac but with only 50% electrode contact area (achieved with using a plastic insulative sheet with a 2.5×2.5 cm annulus); 4) Skin cleaning with alcohol only; 5) Skin cleaning with saline and application of polyvinylpyrrolidone (PVP-K90, 1% w/v) (Sigma-Aldrich Inch, St. Louis, Missouri); 6) Skin cleaning with saline and application of PVP-K90 plus potassium chloride (KCL, 0.0003%). Pretac or PVP-K90 was applied to the disposable electrode surfaces (300-500 uL) before placement on the skin. Electrodes were positioned according to the standard bifrontal placement, avoiding hairline, with careful attention to ensure uniform electrode-skin contact. Static impedance was measured using both a SpECTrum 5000Q (MECTA Corporation, Tualatin, OR) and Thymatron System IV (Somatics LLC) ECT device, immediately after electrode application (t=0) and every minute for 17 minutes or until a stable impedance was recorded. Each subject and electrode preparation combination were tested 3 times, with tests separated by >1 day.

### RUL and BL Clinical ECT Data Set

Anonymized data was re-analyzed from a single-center ECT trial [15] using right unilateral (RUL) and bilateral (BL) ECT. For this series, only stimulation at 800 mA was included in our analysis. Ninety patients in an episode of major depression were randomized to four groups in a 2×2 design, varying both electrode placement and pulse width (0.3 ms vs. 1.5 ms). Except for the pulse width manipulation, stimulation waveform was identical in the ultra-brief and brief pulse groups. 5 cm stainless steel circular disk electrodes were used with hand-held electrode assemblies (MECTA Corporation). Seizure threshold was quantified at the first and last treatments using a titration procedure. At all other treatment sessions electrical dosage was 2.5 or 6.0 times the seizure threshold quantified at the first titration session for BL ECT and RUL ECT, respectively. While precision of seizure threshold may be limited by resolution of titration steps and “floor effects” [8] it is relatively established this approach shows increasing seizure threshold with decreasing dynamic impedance [8,9] for RUL and BL montages.

### BF Clinical ECT Data Set

Anonymized data was analyzed from a single-center ECT trial series using BF ECT. 4.2 by 4.9 cm disposable adhesive electrodes were used (Thymapads, Somatics LLC). High resolution T1-weighted anatomical MRIs were deidentified from a cohort of subjects (n=17) receiving ECT (see modeling below). Each subject received 6-10 ECT sessions with electrodes configured in a bifrontal montage (with Pretac preparation).

### General modeling approach

Skin impedance decreases with increasing electrical current density, to a skin-specific asymptote of ~0.1-0.5 S/m at ~500 uA/cm^2^ [16–18]. However, ECT electrode current density (current divided by electrode area) is >30 mA/cm^2^. Overall head impedance decreases with increasing ECT current density, to a subject-specific asymptote of ~200 Ohm at <800 mA and <200 V [5,11]. At low current, minimum skin resistivity varies substantially with tested-conditions and across microscopic layers, spanning ~1×10^−5^ S/m for epidermal stratum corneum and ~2×10^−4^ to 0.2 S/m for layers of dermis and fat [19,20]. This range can be contrasted with a skull conductivity of ~0.01 S/m.

It is intractable to model <0.1 mm thick tissue layers at full head-scale, necessitating approximation [21]. Adjacent tissue layers with mismatched resistivity can especially impact on current flow patterns [22,23]. Given the above, we consider two scalp layers: a “superficial-scalp” layer and “deep-scalp” layer in the models. The deep-scalp layer is considered to have low conductivity, that is fixed (not effected by current flow). The superficial scalp has minimal conductivity at low current (conditions of static impedance) and high conductivity at high current (conditions of dynamic impedance).

The capacitive effects in biological tissues can be generally neglected when modeling electrical stimulation [24] and are negligible when measured during ECT [10]. Lumped-parameter model of skin use capacitors in parallel with non-linear (current dependent) resistances [16,17]. Capacitive properties (reactance) of tissue, notably skin layers, decrease at either high current [25] or at high frequency. Indeed, the notion that low-intensity (sub-perception) high-frequency (>7 kHz) current can be used during preparation to estimate subject resistance during passage of the ECT stimulus dates to at least 1942 [26], and is reflected in contemporary use of high-frequency to test static impedance. Therefore, we do not model tissue permittivity (reactance). We represent the non-linear changes in resistivity to current flow.

### Subject head segmentation, subject-generic tissue parameterization, and electrodes

Of the seventeen subjects in this clinical series cohort, high resolution MRI-derived head models were constructed for four subjects (#21908, #22615, #22035, #21778), selected based on a range and variance in static and dynamic impedance values. An automated segmentation pipeline based on algorithms in SPM8 [27] and updated for volume conduction models [28] was used to create initial image masks of scalp, skull, air, meninges/cerebrospinal fluid, gray matter and white matter (Fig. 1 B). Additional manual segmentation was applied to correct for noise, aliasing artifacts, and to separate superficial scalp and deep scalp layers.

**Figure 1.**
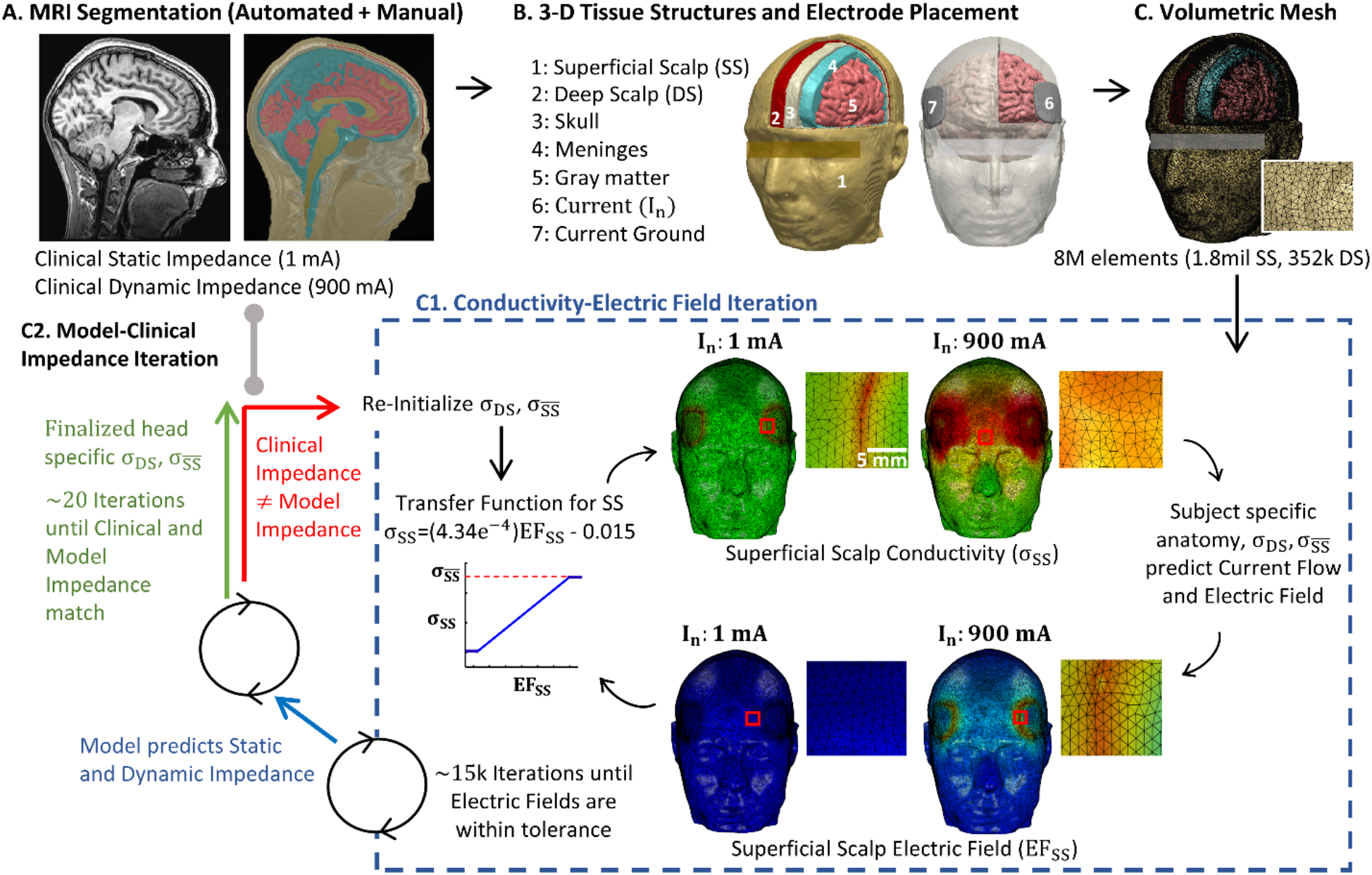
Novel adaptive computational pipeline for ECT current flow FEM models. We developed the first image-derived numerical models of transcranial electrical stimulation (tES) incorporating local changes in tissue conductivity in response to local current flow (electric fields). (A) T1-weighted anatomical MRIs were collected from ECT patients, with static impedance and dynamic impedance data. (B) Volume conductor models were created preserving image resolution using methods previously developed for low-intensity tES [22] [30] [21] - however here we divide scalp into Superficial Scalp (SS) and Deep Scalp (DS) compartments. Skull, meninges, gray matter, and white matter compartments are assigned standard fixed tissue conductivities. Clinical electrode montages are reproduced (e.g. BF) with boundary conditions corresponding to static impedance (I_n_ = 1 mA) and dynamic impedance (I_n_=900 mA) testing. (C) For each subject and electrode montage, a volumetric mesh was generated from the segmented data. (D) The model was initialized with a deep scalp conductivity (σ_DS_) and a maximum superficial scalp conductivity 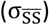. Independently for 1 mA and 900 mA current, an iterative model then computed current flow based on tissue resistivities, updating superficial scalp conductivity 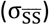 in each element based on local electric fields using a transfer function, and then recalculated brain current flow (blue dashed square). The model converges after ~15k iterations, producing a model prediction of static impedance (for 1 mA) and dynamic impedance (for 900 mA). (E) Model predicted static and dynamic impedance were compared with clinical static and dynamic impedance from the subject. If there was any significant mismatch, the model was reinitialized with updated deep scalp conductivity (σ_DS_) and a maximum superficial scalp conductivity 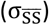, and the FEM was re-run until convergence (blue dashed square). When model static and dynamic impedance matched clinical data, a patient specific deep scalp conductivity (σ_DS_) and a maximum superficial scalp conductivity 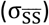 was set.

Unless otherwise indicated, segmented tissues were assigned fixed electrical conductivity [21]: skull (σ = 0.01 S/m), gray matter (σ = 0.276 S/m), white matter (σ = 0.126 S/m), meninges/cerebrospinal fluid (σ = 0.85 S/m), and air (σ = 1e ^−15^ S/m). Deep-scalp layer was assigned a subject specific conductivity (σ _DS_) between 4.5e^−4^ S/m and 0.008 S/m. Local superficial-scalp layer conductivity (σ_SS_) was a function of local scalp electric field (E_SS_) given by:

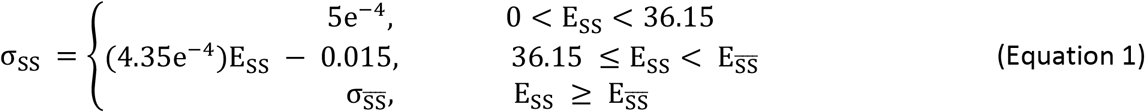

Where 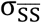 is subject specific maximum superficial-scalp conductivity. We emphasize that E_SS_ varies across the scalp surface such that σ_SS_ then varies across the head (higher near electrodes). For each subject two model parameters were individualized: deep-scalp layer conductivity (σ_DS_) and maximum superficial-layer conductivity 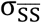.

Ultimately, this is a novel heuristic approach that is computational tractable and support data interoperation and device design (see Discussion). This approach broadly borrows from lumped-parameter skin impedance models that consider a current-sensitive superficial layer and a current-insensitive deep layer with relatively low conductivity [16–18]. A previous model simulating ECT during pregnancy [29] implemented a single skin layer with (un-saturated) conductivity changes restricted under the electrodes.

We modeled two electrode configurations, where the particular protocol used position the electrodes in the corresponding clinical series was reproduced:

1. Bifrontal (BF) pad electrode montage: The center of the both electrodes are on a first imaginary line that starts at the lateral canthus and projects up parallel to the facial midline. The long edge of the electrodes is aligned parallel to the first imaginary line such that the short edge of the pad electrodes is approximately parallel to the horizontal plane. The short edge of the pad electrodes is right above supraorbital ridge (approximately above the eyebrow). Depending on the subject, the center of the electrodes ends up approximately 3-5 cm above the canthus of the eye.
2. Right Unilateral (RUL) disk electrode montage: The center of the frontotemporal disk electrode is aligned with the midpoint of an imaginary line between the tragus and lateral canthus of the eye. The bottom of the disk electrode is placed superior to the imaginary line. The superior electrode disk is centered and to the right of the vertex, which is the intersection of imaginary lines from nasion to inion, and the intertragal line.

Stimulation electrodes and gels were modeled in SolidWorks (Dassault Systèmes Corp., Waltham, MA). For the BF montage, we represented ECT adhesive pad electrodes with dimensions of 4.2 by 4.9 cm and thickness of ~ 1.7 mm, and gel conductivity of 0.018 S/m (based on Thymapad electrodes). For RUL montage we represented circular metal electrodes with a diameter of 5 cm and paste conductivity of 0.018 S/m.

### Computation and subject specific tissue parameterization

These modeled electrodes and gels were incorporated into the segmentation. Volume meshes were generated from the segmented data and exported to COMSOL Multiphysics 5.5 (COMSOL Inc., MA, USA). The resulting mesh comprised > 9,000,000 tetrahedral elements (>15,000,000 degrees of freedom).

The Laplace equation ∇. (σ ∇ V) = 0 (V: scalar electric potential; ∇: gradient vector; σ: conductivity) was solved and the boundary conditions were used such that current in static models (1 mA) and dynamic models 900 mA (unless otherwise stated) is applied to one of the electrode terminals, while the other electrode is grounded. Superficial-scalp conductivity was expressed as a function of electric field (equation 1). The finite element method (FEM) model was implemented using COMSOL. To converge the solution (Fig. 1 C1), a linear system solver of conjugate gradients was used with a relative tolerance of 1×10^−3^ with a nonlinear system solver using the Newton-Raphson method (<500 iterations). This method is applied to millions of degrees of freedom iteratively. Note non-adaptive approaches would require a single Newton-Raphson iteration requiring ~100-500 iteration of the conjugate gradient solver. The total iterations increase to ~2-10k in the adaptive model involving ~60-100 Newton-Raphson iterations.

For each subject (#21908, #22615, #22035, #21778), experimentally reported static impedance and dynamic impedance were determined. Clinical static impedances were averaged across the first non-titrated stimulation of each session, while the dynamic impedances were averaged across the final stimulation of each session (where the seizure was generated). An iterative approach (Fig. 1 C2) was used to search for each subject-specific deep-scalp layer conductivity (σ_DS_) and maximum superficial-layer conductivity 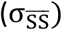 such that model static impedance and dynamic impedance matched subject-specific clinical values.

#### Statistical Analysis

The lilliefors test was conducted to access normality of single-centered ECT trial data comparing montage (RUL vs BL) and waveform (brief pulse vs ultra-brief pulse). It was performed for the static and dynamic impedance data of all subjects that had impedance recordings. A log10 transform was carried out on the nonparametric data and all static and dynamic impedance data was analyzed with a linear regression. Normality of static impedance across electrode preparations in healthy subjects was evaluated with lilliefors test. A Kruskal-Wallis ANOVA determined the statistical significance of the nonparametric data for each instrument, and post-hoc multiple comparison analysis compared means. The relationships between static impedance and dynamic impedance, static impedance and seizure charge threshold, and dynamic impedance and seizure charge threshold of data from the clinical BF ECT series were analyzed using a linear regression after a log10 transform.

## Results

### Relation between static impedance and dynamic impedance in single-center ECT trial

While static impedance and dynamic impedance has long been recognized as measures of individual resistance to electroconvulsive therapy (ECT), their etiology, correlation, and consequences – including how they impact on seizure induction – remain a matter of investigation. Practically, an atypically high static impedance may suggest an undesirably high dynamic impedance, and hence the need to correct electrode setup [31](see below). However, such aberrant impedances may be distinct from less extreme and naturally occurring variation in head impedance (e.g. under ideal electrode site preparation). We conducted a retrospective analysis of static impedance, dynamic impedance, seizure threshold (minimum charge for seizure induction), and patient demographic data recorded from 90 patients (with a total of 622 ECT stimulations) studied in a previously reported clinical trial [15].

ECT data comparing montage electrode placement (RUL vs BL) and stimulus waveform (brief pulse vs ultra-brief pulse) were re-analyzed from this single-center trial [15] – importantly for our purposes, other stimulation parameters were reliably controlled and fixed (e.g. device, electrodes, operator experience, preparation technique). Across ECT conditions, dynamic impedance reliably increased with static impedance (Fig 2) with significant interactions both across subjects within ECT condition and within subject across repeated sessions (not shown). Across stimulation montages and waveforms, there was an evident relationship between static and dynamic impedances (RUL brief: F(2,19)= 9.99, *P* <0.05, R^2^=0.345; BL brief: F(2,38)= 27.5, *P* <0.05, R^2^=0.42; RUL ultra-brief: F(2,21)= 67.3, *P* <0.05, R^2^=0.762; BL ultra-brief: F(2,22)= 33.6, *P* <0.05, R^2^= 0.604). This analysis shows that when a consistent ECT preparation procedures are followed, a correlation between static impedance and dynamic impedance is evident; this is consistent with some aspect of individual anatomy increasing both static and dynamic impedance.

**Figure 2.**
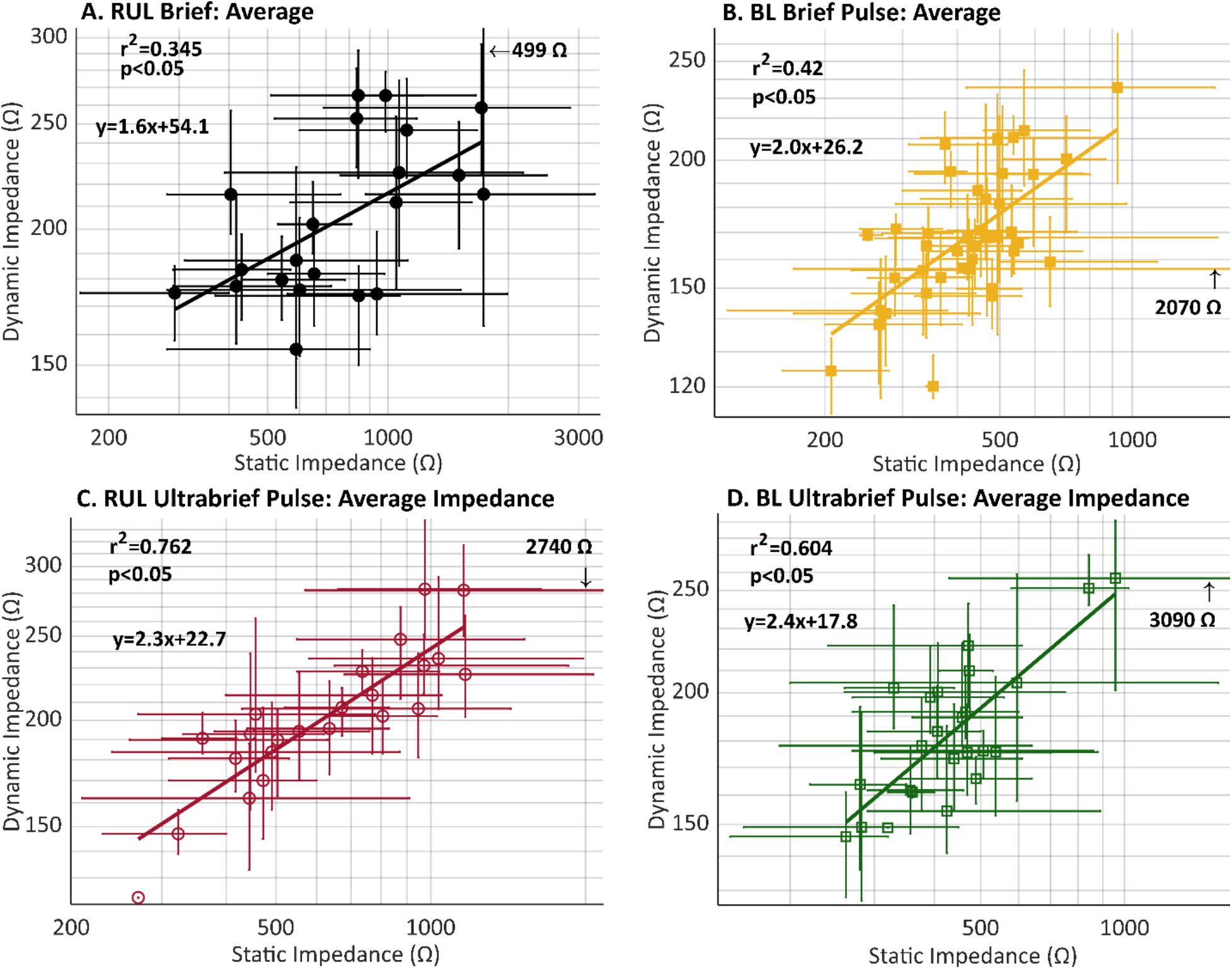
Retrospective analysis of data from a single center ECT trial comparing RUL and BL electrode placements and brief and ultra-brief pulse stimulation. Across (A) RUL Brief Pulse (black), (B) BL Brief Pulse (yellow), (C) RUL Ultrabrief Pulse (red), and BL Ultrabrief pulse (green) there was a significant correlation between patient static impedance and dynamic impedance. The average impedance across sessions is represented for each subject as well as the full range of variance (error bars).

### Impact of electrode preparation technique on static impedance

In a sample of healthy subjects, we measured static impedance across systematically varied adhesive electrode preparation techniques (Fig. 3). Each condition was tested repeatedly across subjects (n=3) and the changes over time were monitored. All electrode preparation conditions using an adhesive solution applied to the electrode surface (Pretac or PVP-K90) resulted in a comparable and minimal static impedance for each given subject (Thymatron: *χ*^*2*^ *=* 59.82, df = 5, P < 0.05; spECTrum: *χ*^*2*^ *=* 63.75, df = 5, P < 0.05). Static impedance increased in the absence of the adhesive (alcohol skin cleaning only). Reduction in electrode contact area by 50% increased static impedance a factor of ~1.5-2.7 (Fig.3). Excluding conditions of limited electrode-skin contact, the lower static impedance reported by the MECTA compared to the Somatics device did not impact rank order or reliability.

**Figure 3:**
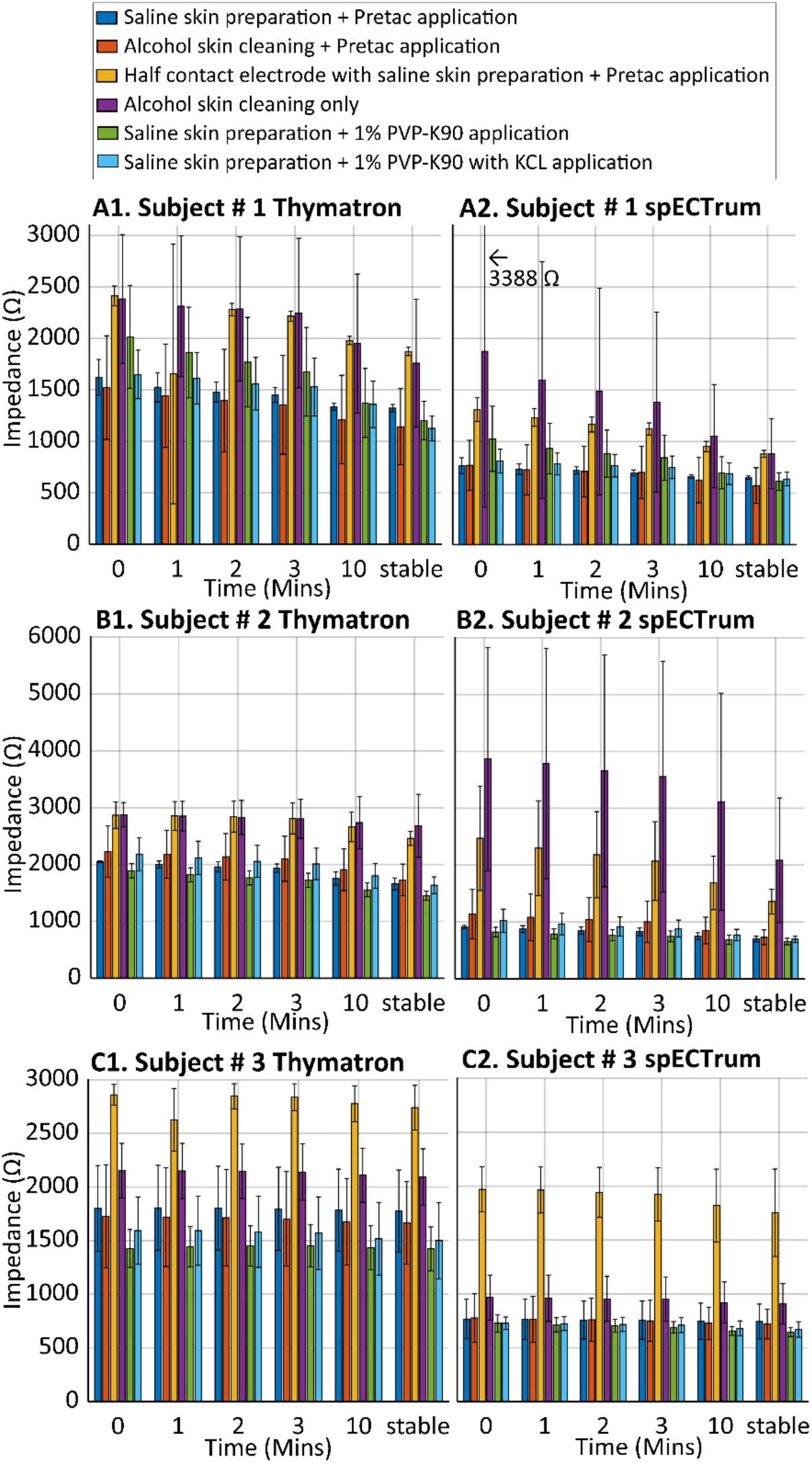
Static impedance over time under six electrode preparations. Each condition measured at least three times on the Thymatron (1) and SpECTrum (2) devices in subject 1 (A), subject 2 (B), and subject 3 (C).

Even under our tightly controlled experimental conditions and with optimal preparation (use of adhesive), moderate variability was observed between repetition trials. Under realistic clinical conditions, variability both in the quantity (area coverage) and quality (e.g. extent of adhesion) of electrodes could produce substantial impedance variance. Nevertheless, it is important for our modeling efforts that, assuming good ECT electrode preparation, difference between individuals are not dominated by idiosyncrasy in the quality of electrode-skin contact (i.e. as models assume complete electrode-skin contact) but rather reflects subject’s head properties.

### Development of individualized adaptive tissue models of ECT

Previously MRI-derived head models to predict subject-specific brain current flow were developed [22,32] and validated [33,34] for low-intensity tES, such as tDCS, and then subsequently applied to ECT [14,35]. These models assumed non-adaptive tissue resistivity. Here we applied to ECT data our novel pipeline (Fig. 1) for adaptive tissue conductivity (where local conductivity changes with electric field strength (Fig. 4). The analysis was conducted on data contributed by four ECT patients. Each patient’s anatomical MRI was segmented and an iterative search process identified the maximum superficial scalp conductivity 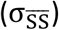 and the deep scalp conductivity (σ_DS_) that produced a prediction of static and dynamic impedance values corresponding to the patient’s clinical data. The resulting subject specific parameters were: Subject 21908, 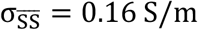 at 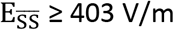, σ_DS_ = 0.002 S/m; Subject 22615, 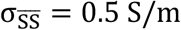 at 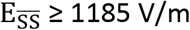, σ_DS_ = 4.5e − 4 S/m; Subject 22035, 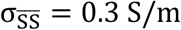 at 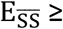 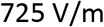, σ_DS_ = 0.008 S/m; Subject 21778 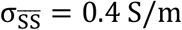, at 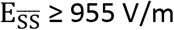, σ_DS_ = 0.0012 S/m.

**Figure 4.**
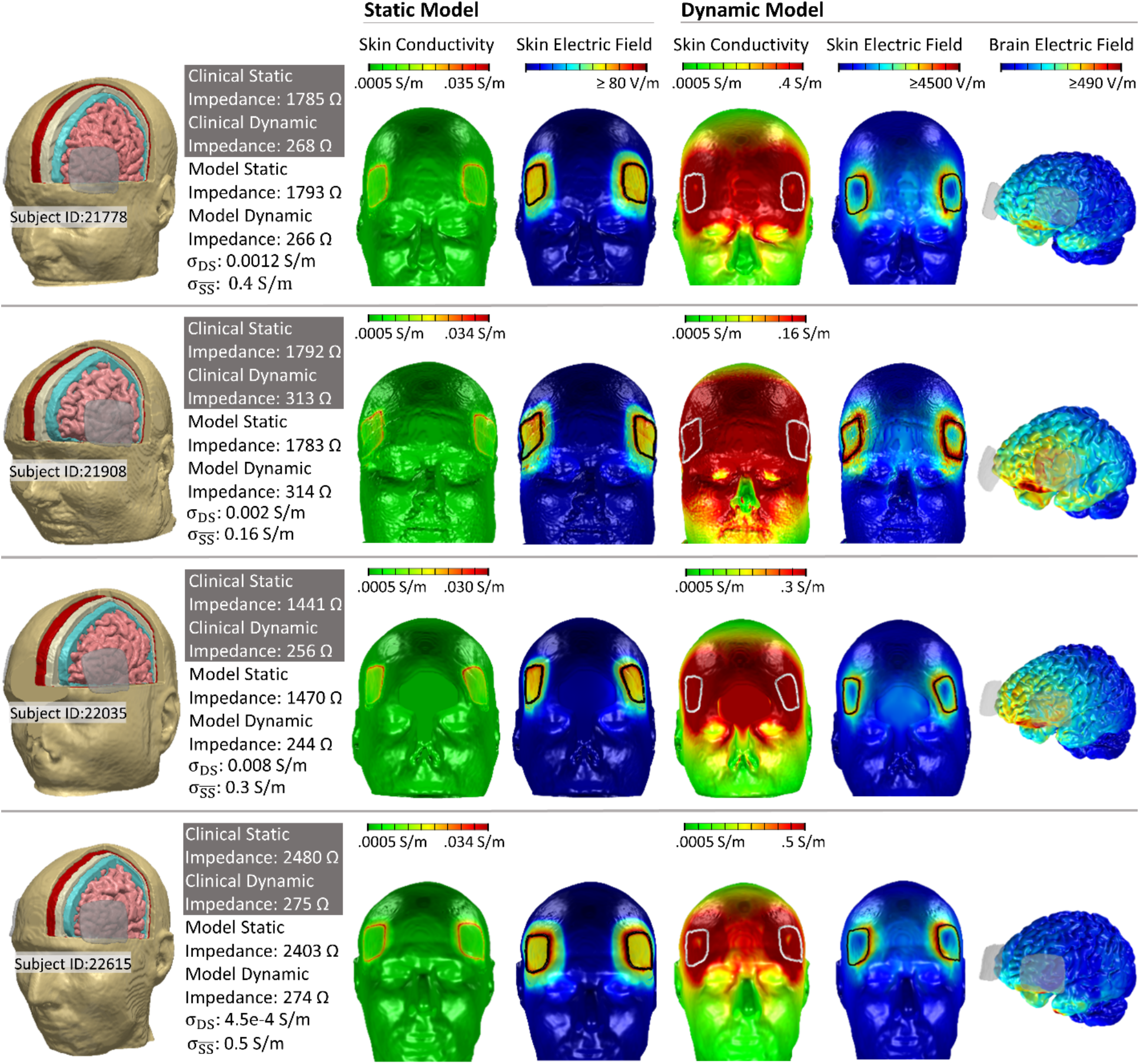
Dynamic ECT models for four ECT subjects. Dynamic FEM models simulated current flow across four subjects who have received ECT (Subject IDs: 21778, 21908. 22035, 22615). (First Column) Models anatomy was based in subject anatomical MRI. Static impedance and dynamic impedance values were recorded for each subject (grey box). Each subject model was assigned a specific deep scalp conductivity (σ_DS_) and a maximum superficial scalp conductivity 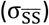 as indicated, such that adaptive FEM simulation predicted corresponding static impedance (based on 1 mA applied current) and dynamic impedance (based on 900 mA applied current) as indicated. (Second and Third Column) Results from the static impedance (1 mA current) simulation showing resulting superficial scalp conductivity and scalp electric field. (Third, Fourth, Fifth Column) Results from the dynamic impedance (900 mA current) simulation showing resulting superficial scalp conductivity, scalp electric field, and brain electric field. We emphasize in these novel adaptive simulations that brain current flow was determined by tissue conductivity, superficial scalp conductivity was simultaneously determined by local electric field. Even for the 1 mA (static) model local changes in scalp conductivity are predicted. For the 900 mA (dynamic) model, the saturation of the transfer function between superficial scalp electric field and conductivity results in a more diffuse saturation of scalp conductivity (front of head) compared to scalp electric field (around electrodes).

Stimulation with 1 mA (Fig. 1, static model) produced peak electric fields under and around electrode edges (>80 V/m) with moderate increases in conductivity (~0.03 S/m) around the electrode perimeters. Stimulation with 900 mA produced high electric field across the scalp forehead with peaks around electrodes (>45000 V/m), and an associated increase in scalp conductivity (0.15-0.5 S/m). The resulting brain current flow during ECT also predicted peak electric fields >490 V/m.

### Adaptive scalp response and the relation between static and dynamic impedance

The role of head anatomy in ECT outcomes remains unclear, and is complex to understand experimentally when gross anatomy, tissue properties, and neurophysiology all vary across individuals. The role of gross anatomy can be considered using computational models by 1) fixing head anatomy and manipulating tissue (scalp) properties; or 2) artificially changing (e.g. dilating) anatomy in a single head. These approaches are used first to explain the relation between dynamic and static impedance, and then how these impact delivery of current to the brain.

For each head we fitted deep-layer scalp conductivity (σ_DS_) and maximum superficial-layer scalp conductivity 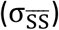. Subsequently, it is possible to examine the role of scalp properties by simulating the “swapping” of the scalp properties across different heads (Fig. 5). Starting from a base set of four head anatomies (Subject 22615, square; Subject 21778, diamond; Subject 22035, circle; Subject 21908, x with corresponding σ_DS_ and 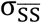 for each subject), we then swapped (mixed) scalp properties across heads (colors), resulting in 12 synthesized heads (plus the 4 originals). Under an assumption that a theoretical subjects head anatomy, deep-layer scalp conductivity (σ_DS_) and maximum superficial-layer scalp conductivity 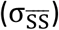 can vary independently, each synthesized head represented a novel hypothetical subject. This approach allows systematic comparison of the relative impact of anatomy and scalp properties to ECT outcomes as predicted by the synthetic models.

**Figure 5.**
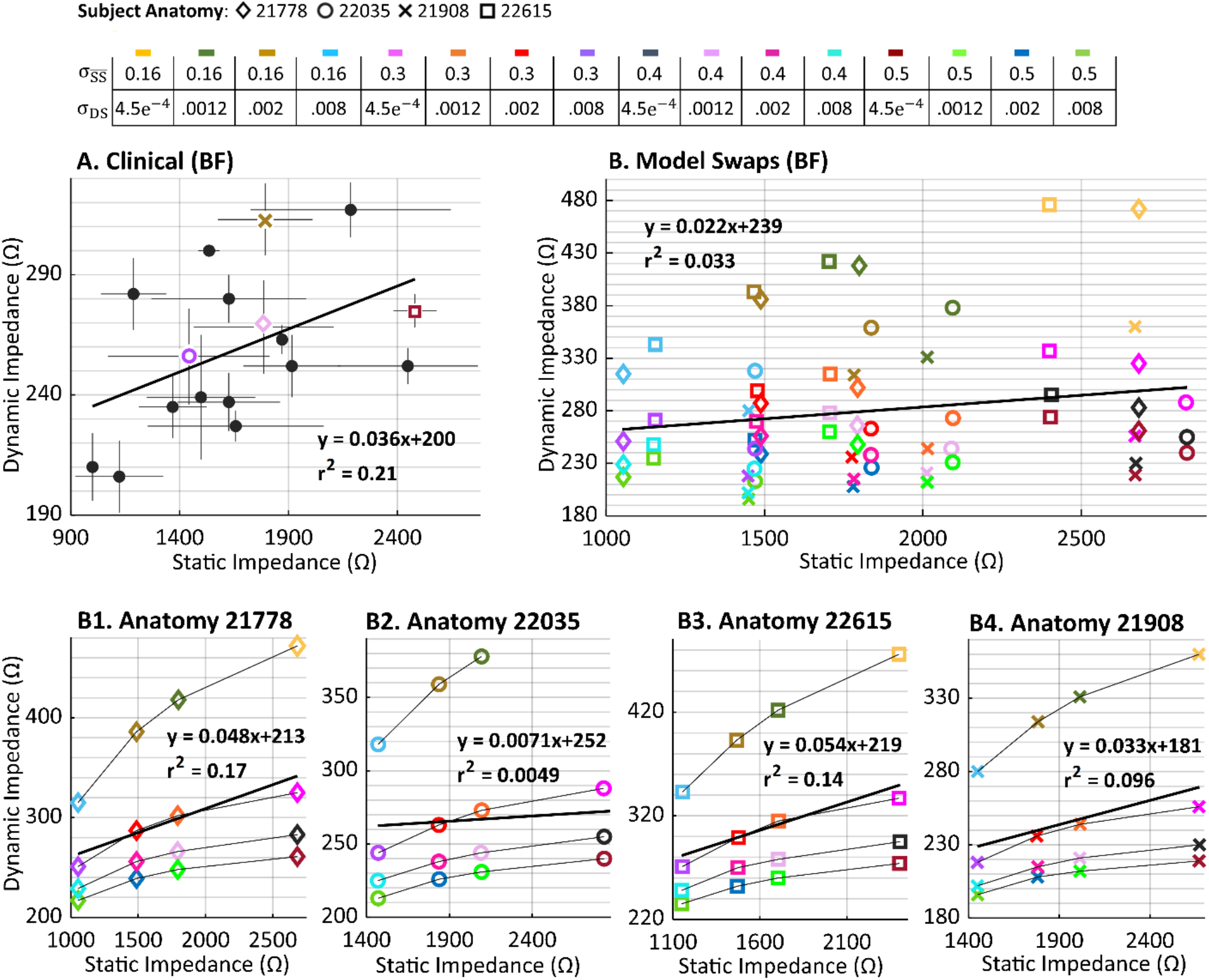
Relation between Dynamic Impedance and Static Impedance in clinical data and adaptive models including scalp swaps. (A) Dynamic impedance and static impedance were modestly correlated in an exemplary patient sample receiving BF ECT. Four subjects (colored symbols) were selected for adaptive FEM simulation. (B) In these four subjects, static and dynamic impedance values were simulated by assigning subject specific deep scalp conductivity (σ_DS_) and a maximum superficial scalp conductivity 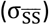. The conductivities (in S/m) are called “endogenous” for our purposes here (Subject 22035 σ_DS_= 0.008, 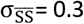; Subject 21778 σ_DS_= 0.0012, 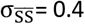; Subject 21908 σ_DS_= 0.002, 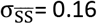; Subject 22615 σ_DS_= 4.5e^−4^, 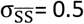). In scalp swaps, all deep scalp conductivity (σ_DS_) and a maximum superficial scalp conductivity 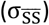 were varied across all subject anatomies, resulting in 16 simulates swapped-conductivity heads (including the endogenous 4 heads). For each head, static impedance and dynamic impedance was predicted using our adaptive pipeline. There was only a weak correlation between simulated static and dynamic impedances. (B1, B2, B3, B4) Replotting the same points but separated by subjects, highlighted certain trends. For a given head anatomy and a given deep scalp conductivity (σ_DS_), varying superficial scalp conductivity 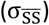 changes dynamic impedance but not static impedance. This vertical distribution of dynamic impedances for a given static impedance reduces overall correlation across all simulations, but even within a single anatomy. For a given head anatomy and superficial scalp conductivity 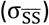, decreasing superficial scalp conductivity 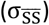 monotonically increases both static impedance and dynamic impedances, explaining the source of overall (weak) correlation.

The relationship between static impedance and dynamic impedance is evident though clearly imperfect in the data from the clinical BF ECT series (Fig. 5 A). With dynamic impedance as the dependent factor, the group yielded statistically significant results (Clinical BF: F(2,15)= 5.64, *P*<0.05, R^2^=0.273). A consentient relationship between static impedance and dynamic impedance across all the synthetic heads was not evident (Fig 5 B) though interactions are evident when considering individual head anatomy (Fig. 5 B1, B2, B3, B4).

Simulations with swapped-scalps predict variation in scalp properties are relatively more important than gross anatomy in determining static impedance and dynamic impedance. Specifically, extreme variations in static impedance and dynamic impedance are governed by 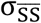 and σ_DS_ respectively. Therefore, because 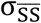 and σ_DS_ vary independently a dispersion of static and dynamic conductivities is produced. A weak interaction persists because σ_DS_ has (in addition to a strong effect on static impedance) a weak effect on dynamic impedance (Fig 4 B1, B2, B3, B4 lines represent fixed head and 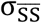, so only σ_DS_ varying). Head anatomy moderate both static and dynamic impedance (see also below). The importance of deep-layer scalp (σ_DS_) conductance, with head anatomy as a moderator, on static impedance is evident when noting that for a given head and σ_DS_,static conductivity is nearly fixed, despite variation in dynamic impedance. This vertical dispersion is also why a relationship between static and dynamic impedance is not evident across swapped-scalp models (e.g. for static impedance >2600 ohm a wide range of dynamic impedance is possible).

Across (almost all) variations in tissue properties, the relative (rank) order of dynamic impedance was fixed (from higher to lowest: Subject 22615, square; Subject 21778, diamond; Subject 22035, circle; Subject 21908, cross). Across variation in adaptive-tissue properties, the rank order of static impedance was not consistent across heads, though some trends were evident. Subject 22035 presented the highest relative static impedance. Subject 22615 presented the lowest relative static impedance. In any case, neither rank order for static impedance or dynamic impedance corresponded to the rank order of any global anatomical feature (head circumference, skull thickness, inter-electrode distance). This suggested that it is not possible to predict (relative) static or dynamic impedance based on any simple anatomical measure (consistent with clinical observations; [5]).

The salient result here is not that our adaptive modeling approach can match the clinical impedance data, since adjusting subject 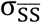 and σ_DS_ (Fig. 4) could ensure model approximation of subject-specific clinical data. Rather, these models propose an explanation for clinical observations debated over decades.

### Adaptive scalp response and the relation between brain current intensity and static or dynamic impedance

In the BF ECT clinical series, there was no evident relation between static impedance and seizure threshold (Fig. 6 B1) and a modest relationship between dynamic impedance and seizure charge threshold (Fig. 6 B2). With seizure charge threshold as the dependent factor, the clinical groups were not statistically significant at *P* <0.05 (A1. Clinical BF: F(2,14) = 0.84, *P* = 0.375, R^2^ = 0.0566; A2. Clinical BF: F(2,10)=1.96, *P* =0.192, R^2^ = 0.164). Across the swapped BF head model stimulations (4 originals plus 12 synthesized scalp heads), we predicted peak electric fields, both brain-wide and specifically in the motor strip. There was no evident correlation between static or dynamic impedance, and brain peak electric field or motor strip electric field. The exception between a weak negative relation between static impedance and motor strip peak electric field.

**Figure 6.**
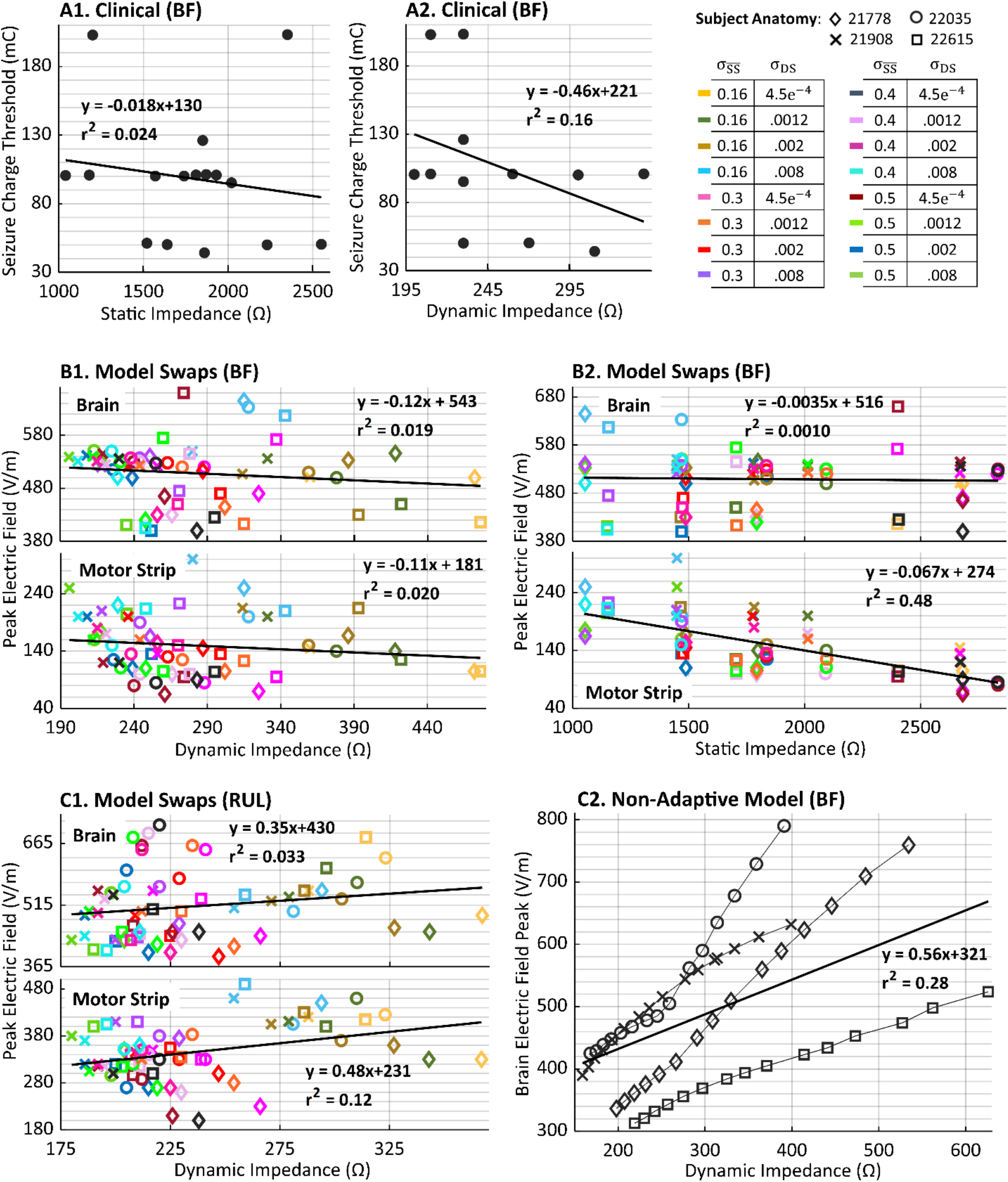
Relation between brain electric field (or seizure threshold) in clinical data and adaptive scalp models including scalp swaps. In theory, conditions that result is higher electric fields (at the region of seizure initiation) will be associated with reduced charge required to trigger a seizure, i.e. lower seizure threshold. (A) In an exemplary patient sample receiving BF ECT, the relation between increasing static impedance (A1) or increasing dynamic impedance (A2) and decreasing charge threshold for seizure initiation. (B) The predicted relation between brain electric field and dynamic impedance (B1) or static impedance (B2) can be simulated for the BF electrode montage using adaptive FEM models of the 16 swapped-conductivity heads (including the endogenous 4 heads). Brain wide peak electric field is considered (higher panel) as well as motor strip peak electric field (lower panel). Only weak correlations are predicted, with the strongest correlation (negative direction) for the case of static impedance and motor strip peak electric fields. (C1) Using adaptive FEM modeling, the relation between dynamic impedance and static impedance was further predicted in same heads for the RUL electrode montage. Weak correlation was predicted, with the strongest relationship (positive direction) for dynamic impedance and motor strip peak electric field. (C2) Non-conventional FEM of the four anatomical head, where for each uniform scalp conductivity was varied to adjust dynamic impedance, predicted quasi-linear relationship for each head between dynamic impedance and brain electric field. Variability between heads resulted in a weaker group correlation between dynamic impedance and brain electric field. We note the prediction in non-adaptive conventional stimulation of increasing electric field with increased dynamic impedance is expected and consistent with scalp resistivity increasing both. The predictions from adaptive FEM suggest a more complex and nuanced relationship between dynamic impedance and electric field (and so charge needed to produce seizures).

While these adaptive models were parametrized based on BF ECT data, we used the same swapped heads to predict for RUL ECT the relation between static impedance or dynamic impedance and brain current delivery (Fig. 6 C1). Relationships were weak, and trended positive. We further considered non-adaptive models in order to highlight the unique behavior of adaptive models (Fig 6 C2). In non-adaptive models, for each subject head, the (uniform) scalp conductivity was incrementally decreased, producing a range of impedance (note, in these non-adaptive models we distinguish static impedance or dynamic impedance). As expected, in non-adaptive models, brain electric field increased monotonically for each subject with increased impedance. This slope varies across heads but remains significant across the group.

**Table 1:** Tissue dilation in adaptive FEM stimulation of four ECT subjects. Simulation results showing static/dynamic impedance along with brain/motor electric field from adaptive computational simulations for each endogenous head before and after tissue dilations. These values are shown for each head’s standard (original) anatomy (third column), dilation of superficial scalp 6-fold (fourth column), and further dilation deep scalp 2-fold (last column). Electric field values from dynamic models and reported as overall peaks in the brain as well as the motor cortex as a region of interest.

|  | Subject ID | Standard Anatomy | Superficial Scalp Dilation 6x | Superficial Scalp Dilation 6x + Deep Scalp 2x |
| --- | --- | --- | --- | --- |
| Static/Dynamic Impedance | #22615 | 2403/274 ($\Omega$) | 2931/206 ($\Omega$) | 3820/219 ($\Omega$) |
| | #21778 | 1793/266 ($\Omega$) | 2428/199 ($\Omega$) | 2944/217 ($\Omega$) |
| | #21908 | 1783/314 ($\Omega$) | 2453/262 ($\Omega$) | 2742/295 ($\Omega$) |
| | #22035 | 1440/244 ($\Omega$) | 2151/199 ($\Omega$) | 2083/216 ($\Omega$) |
| Brain/Motor Electric Field for Dynamic Models | #22615 | 660/95 (V/m) | 411/68 (V/m) | 463/64 (V/m) |
|  | #21778 | 460/115 (V/m) | 320/85 (V/m) | 300/70 (V/m) |
|  | #21908 | 507/215 (V/m) | 438/150 (V/m) | 413/120 (V/m) |
|  | #22035 | 470/160 (V/m) | 400/140 (V/m) | 380/115 (V/m) |

MRI-derived head models can be artificially altered [36] including dilation of specific tissue layers [23,30]. Keeping tissue conductivity properties fixed (as optimized for each subject), we first dilated only superficial scalp layer by ~6x (Fig 7). Superficial-scalp dilation increased static impedance, decreased dynamic impedance, and decreased brain current delivery in all four heads. Further dilation of deep scalp layer by ~2x increased static impedance in three of the four heads, increased dynamic impedance in all four heads, and decreased current delivery in three of the four heads in comparison with the results from superficial-scalp dilation. Notably, these simulations show a dissociation among static impedance, dynamic impedance, and electric field in the sense that specific changes can affect them relatively differently. This also reinforces the unique outcomes, and so value, of adaptive-resistivity models.

**Figure 7:**
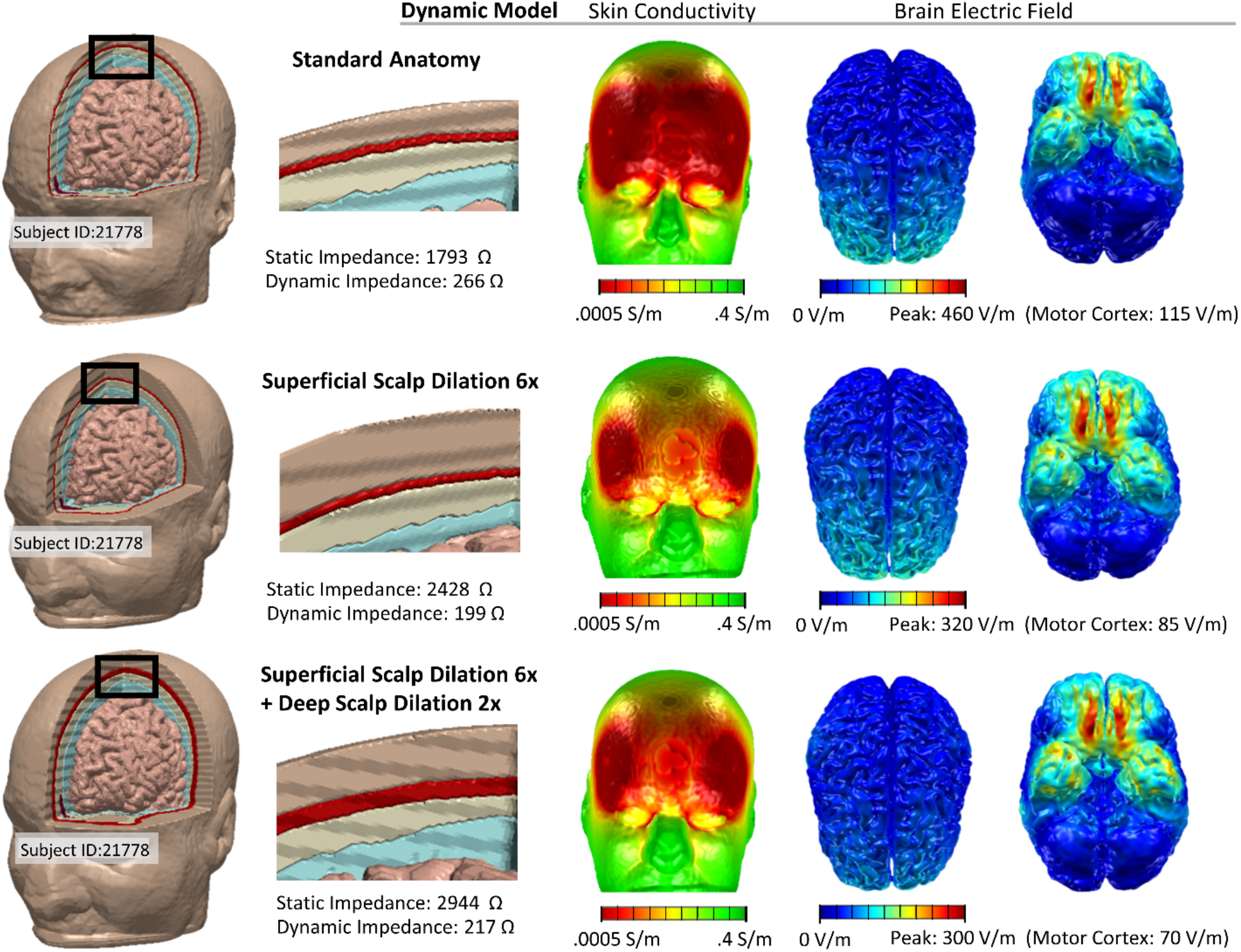
Tissue Dilation in Adaptive FEM model of ECT, and relationship between static impedance, dynamic impedance, and brain electric field. Our adaptive FEM models suggest a complex relationship among head anatomy, tissue properties, and resulting current flow and resistance. In addition to allowing variance of just select tissue conductivity, computational models can access the role of isolated anatomy, including through superficial tissue dilation [23]. We considered standard (original) anatomy (top row), dilating superficial scalp 6-fold (middle row), and further dilating deep scalp 2-fold (bottom row). In the last case, dilated deep scalp replaces overlying superficial scalp. Compared to the standard anatomy, dilating superficial scalp 6-fold increased static impedance, decreased dynamic impedance, and decreased brain electric field. Further dilating deep scalp 2-fold, increase static impedance, increase dynamic impedance, and decreased further brain electric field. These adaptive FEM simulations show a subtle non-monotonic relationship between static impedance, dynamic impedance, and brain electric fields.

### Response to moderated ECT Voltage and Current: Model validation against classical clinical data

While modern ECT uses currents of 800-900 mA, in two earlier trials the current and/or voltage of ECT was systematically varied within subjects, which can also be modelled with adaptive-conductivity simulations (Fig. 8). Dynamic impedance as a function of voltage from Umlauf et al. 1951 and Maxwell et al. 1968, and dynamic impedance as a function of current for Umlauf et al. 1951 were replotted for each individual subject (grey lines) and group average (yellow Maxwell et al. 1968; black Umlauf et al. 1951). Group data are fit by a power law:

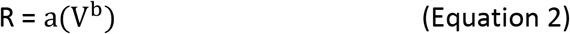

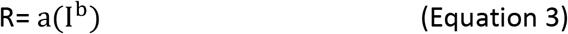

Where R is Dynamic Impedance (in Ohms), V is voltage applied (in Volts), I is current applied (in mA) and a and b are constants.

**Figure 8:**
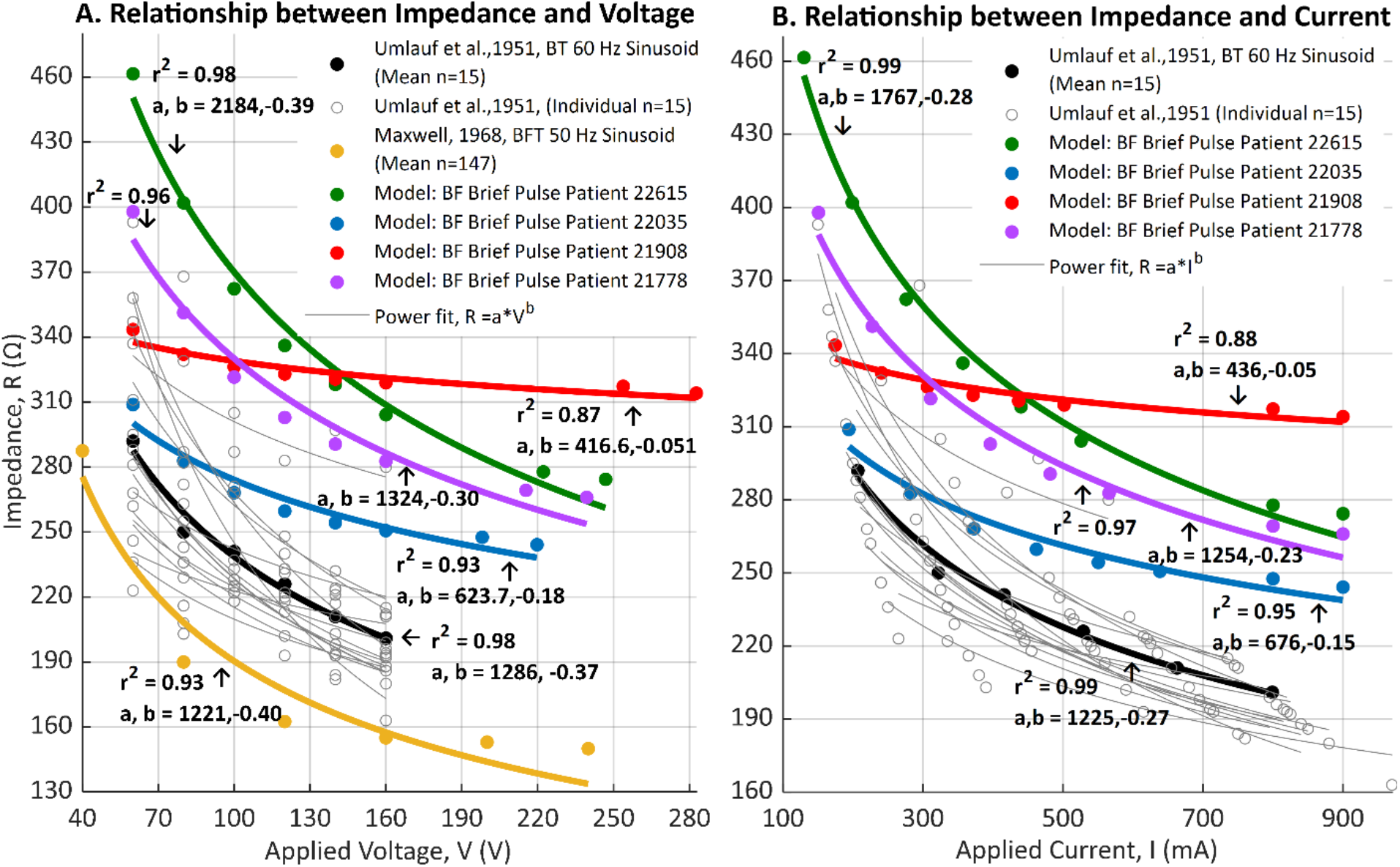
Relationships between varied applied voltage or current in ECT and measured dynamic Impedance: clinical and modeling results. While in conventional ECT the current level if fixed, in historical experiments the current and/or the voltage was reduced while monitoring dynamic impedance. In Umlauf et al. (1951), ECT dosage (using 60 Hz sinusoidal waveforms, BT electrode montage) was varied, with resulting impedance reported as a function of both applied voltage (A) and applied current (B) [5]. Individual subject data (grey) is reported, alongside group average (black). In Maxwell et al. (1968), the voltage of ECT (using 50 Hz sinusoidal waveform, BFT electrode montage) was reported against dynamic impedance (group average only, yellow)[11]. Using adaptive FEM, for each head (Subject IDs: 21778, 21908, 22035, 22615), we systematically varied the current intensity applied in the model and simulated the resulting dynamic impedance (B) while also reporting the associated voltage (A). Our models were parameterized based on pulsed waveforms (only for 1 mA and 900 mA). Average clinical data and individual model data were fit by a power law. While absolute difference between clinical cases and models are expected (e.g. given variation in protocol and the dependence of conductivity on waveform), the power law fits well the clinical and modeling data.

Alongside these clinical data, results from reducing the applied voltage and current in our adaptive-conductivity models heads is shown and fit to power law. The models broadly reproduce the power law relationship. Absolute matching of parameters (a, b) is not expected given different in protocols. Umlauf et al. 1951 and Maxwell et al. 1968 used sinusoidal stimulation and our models parameterized based on rectangular pulse trains. It is known that impedance will vary with pulse/sinusoidal waveforms [37].

## Discussion

Current passage through the scalp (skin) depends on numerous layers and ultra-structures, each with a complex (non-linear, time-dependent) impedance to current flow [16,38–42] – which is computationally intractable for tES head models [21]. The pipeline developed here to simulate adaptive-scalp conductivity during tES represent two scalp layers, with just two respective associated subject-specific parameters: a deep-scalp layer with fixed conductivity (σ_DS_) and a superficial-scalp layer where electric fields increase conductivity up to a maximum 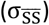. Ongoing modifications and refinements of our pipeline are welcome but, as discussed below, it provides a heuristic solution supporting rational tES/ECT optimization. In this same vein, additional complexity (e.g. adding microscopic sweat ducts [19,38]) may not change the relevant clinical parameters predicted by these models.

The physical properties of the ECT stimulus markedly effect both efficacy and cognitive side effects. It is the combination of ECT dose (electrode montage, waveform, and stimulus intensity) and head resistive features, that determine how much and where current is delivered to the brain. That head resistivity varies across subjects has been known for decades based on clinical measurement of static impedance and dynamic impedance. But it has been hard to explain if and how these head resistances impact on stimulation during ECT (brain current delivery) because their etiology are unclear. The ongoing universal reliance on static impedance and dynamic impedance in ECT stems largely from limitations built into ECT devices (e.g. voltage compliance) and clinical standards (e.g. preparation quality control) [10] and thus consider impedance extreme. Nonetheless, there has been discussion spanning decades on whether less extreme variation in impedance parameters (deriving from endogenous difference in anatomy) can be better leveraged to understand and optimize ECT dosing and behavioral outcomes. Our adaptive-conductivity ECT models make a range of predictions on these matters.

Supporting our analysis, we consider data from ECT treatments using a range of devices, electrode placement, and waveforms (Fig. 2, Fig. 6) showing a definitive but imperfect relationship between static and dynamic impedance - consistent with endogenous differences between subjects increase static and dynamic impedance together. We also show static impedance is impacted by electrode preparation technique (including both electrode-skin contact area and adhesion quality), but endogenous individual difference remains (Fig. 3).

Adaptive ECT models predict that individual difference in static and dynamic impedance largely reflect difference in individual scalp properties, with head anatomy playing a moderating role. A component of individual scalp conductivity that is insensitive to current passage (σ_DS_) governs static impedance (to low currents). Though limited scalp conductivity changes are predicted even at low current (Fig. 4), they do not meaningfully impact static impedance (Fig. 5). Dynamic impedance (to high current) is largely governed by the individual’s maximum adaptive scalp conductivity 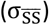 in response to current passage, with a smaller influence of the current-insensitive scalp impedance (σ_DS_). The dual impact of individual σ_DS_ may explain an intrinsic but imperfect relationship between static impedance and dynamic impedance. Contrariwise, the insensitivity of static impedance to the scalp adaption to current flow 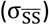 explains the weakness of any relationship between static and dynamic impedance. Namely, variations in just 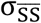 result in the same static impedance being associated with a wide range of possible dynamic impedance. If a given clinical trial reports a correlation between static and dynamic impedance (Fig. 1) or not [5] can reflect difference in electrode / skin preparation or stimulation current. Protocol (e.g. montage, waveform) and subject (e.g. age, sex) differences in impedance would also depend on these scalp properties [9,43].

While static and dynamic impedance are ubiquitous ECT biomarkers supporting model validation (Fig. 8), tES current flow models have translational value in informing treatment protocols [1,29,35,44–46]. These efforts consider the electric fields generated across the brain relevant for both seizure threshold and side-effects. While non-adaptive models predict a direct relation between impedance and brain electric field intensity (Fig. 6 C2), adaptive models suggest a complex relationship (Fig 6 B). Indeed, adaptive ECT models predict a non-monotonic relationship between static impedance, dynamic impedance, and brain currents (Fig. 7). Nevertheless, understanding how adapting scalp properties impact (unmeasurable) brain current flow and how they are reflected in (measurable) impedance parameters, may support efforts to optimize ECT therapy.

## Notes

**Conflict of Interest** The City University of New York (CUNY) has IP on neuro-stimulation systems and methods with authors NK and MB as inventors. MB has equity in Soterix Medical. MB served on the advisory boards, received grants, and/or consulted for Boston Scientific, MECTA Corporation, Halo Neuroscience, Biovisics, and GlaxoSmithKline Inc. HAS serves as a scientific adviser to Cerebral Therapeutics, LivaNova PLC, MECTA Corporation, and Neuronetics Inc. He receives honoraria and royalties from Elsevier, Inc. and Oxford University Press. HAS is the inventor on non-remunerative US patents for Focal Electrically-Administered Seizure Therapy (FEAST), titration in the current domain in ECT, and the adjustment of current in ECT devices, each held by the MECTA Corporation. HAS is also the originator of magnetic seizure therapy (MST).

### Competing Interest Statement

The authors have declared no competing interest.

